# *Listeria monocytogenes* faecal carriage is common and driven by microbiota

**DOI:** 10.1101/2021.01.13.426560

**Authors:** Marc Garcia-Garcera, Lukas Hafner, Christophe Burucoa, Alexandra Moura, Maxime Pichon, Marc Lecuit

## Abstract

*Listeria* genus comprises two opportunistic pathogenic species, *L. monocytogenes* (*Lm*) and *L. ivanovii*, and several non-pathogenic species. All can thrive as saprophytes, whereas only pathogenic species cause systemic infections in human and cattle. Identifying *Listeria* species’ respective biotopes is critical to understand the ecological contribution of *Listeria* pathogenic potential. Here, we aimed at detecting *Listeria* in samples of diverse origins, to highlight ecological differences between pathogenic and non-pathogenic species. We retrieved 16S rDNA datasets from the metagenomics MG-RAST database and determined the prevalence and abundance of *Listeria* species in various sources. Overall, *Listeria* was detected in 14% of datasets. *Lm* was the most prevalent species, most abundant both in soil and host-associated environments, including in 5% of human stools. *Lm* was also detected in 10% of human stool samples from an independent cohort of 900 healthy asymptomatic donors. A specific microbiota signature was associated with *Lm* faecal carriage in human, as well as in experimentally inoculated mice, in which it preceded *Lm* long-term gut colonization, indicating that gut microbiota composition influences *Lm* faecal carriage. These results suggest that asymptomatic faecal carriage, rather than disease, exerts purifying selection on *Lm* “virulence genes”.

## Introduction

Infectious disease symptoms can favor the transmission of pathogenic microorganisms and hence select for genes that induce these symptoms (e. g. cough induced by *Mycobacterium tuberculosis*^1^). However, asymptomatic host colonization can also favor microbial transmission, and thereby select for genes involved in host-microbe association that may also be involved in the development of opportunistic infections. *Listeria monocytogenes* (*Lm*) and *L. ivanovii* can cause opportunistic infection in human and other mammals including cattle^2,3^, leading to fetal-placental infection, abortion and encephalitis, in contrast to other *Listeria* species which are non-pathogenic. *Lm* is known to alternate between a saprophytic and a host-associated lifestyle during which it expresses so-called virulence factors that mediate tissue invasion and within-host dissemination^4^. Most of these virulence factors are part of *Lm* core genome and subjected to purifying selection^5–7^. *Lm* most virulent clones are also the most adapted to mammalian gut colonization^8^ and *Lm* can be released from infected tissues back to the intestinal lumen^6,9,10^, indicating that virulence may ultimately promote *Lm* faecal carriage and thereby play a major role in its dissemination.

*Lm* is a common contaminant of foodstuffs, and each human individual in Western countries is estimated to be exposed to *Lm* multiple times per year^11^. Yet the incidence of microbiologically proven invasive human listeriosis is extremely low, with 0.28 and 0.6 cases *per* 100,000 people in the US and Europe, respectively^12,13^. This implies that in most cases, human exposure to *Lm* leads to either absence of infection, and/or clinically silent gut colonisation, and suggests that *Lm* virulence genes are likely not selected for their capacity to induce clinically overt disease. There have been reports of *Lm* asymptomatic faecal carriage, both in human and cattle^14–23^, and all large scale studies have suggested that the prevalence of *Lm* carriage is below 1%^18–20^. However, these studies were based on culture-based methods^18–20^, which are less sensitive when directly compared to molecular detection methods like PCR and sequencing^21,22,24^. Large molecular studies on the distribution of *Listeria* species in mammals and the environment are not available^25–27^.

## Results

Ecological sampling is influenced by *a priori* assumptions about potential niches^28,29^. Here we circumvented this limitation by assessing *Listeria* species distribution in publicly available metagenomic datasets from the large MG-RAST database^30^, to which high quality metadata are associated, and retrieved 2,490 full metagenomes and 11,907 16S rDNA high quality datasets (see Materials and methods). We assessed the impact of *Listeria* pathogenic potential on its ecological distribution by comparing the relative abundance (proportion of a species in a given sample, x axis, Fig. 1) and prevalence (frequency of a species in samples of a given category, y axis, Fig. 1) of the *Listeria* pathogenic species *Lm* and *L. ivanovii* to that of the non-pathogenic species *L. innocua*, *L. seeligeri* and *L. welshimeri*^31^.

**Figure 1.**
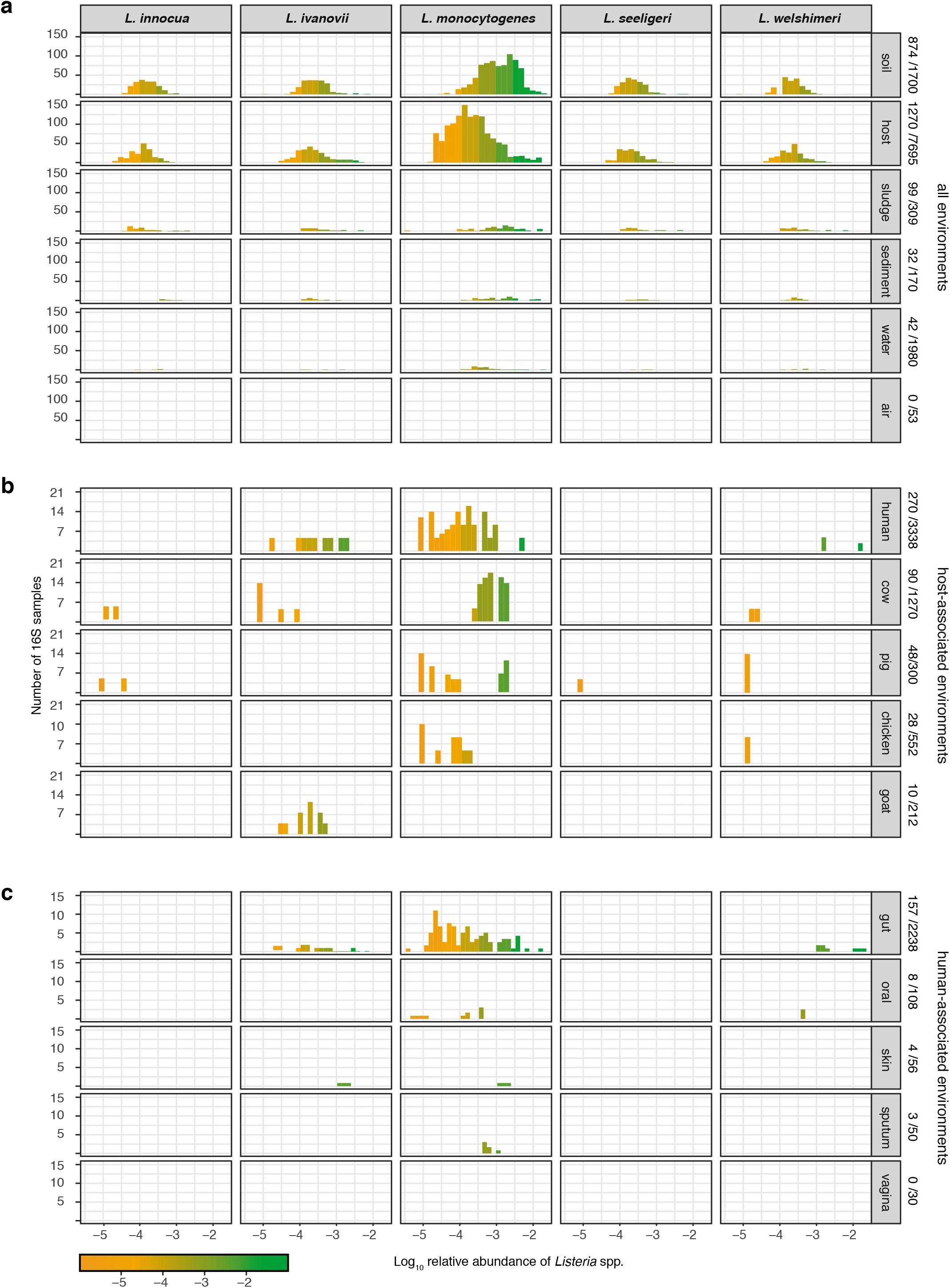
*Lm* is more prevalent in host-associated environments than non-pathogenic *Listeria* species. Relative abundance and prevalence of *Listeria sensu stricto* species in 16S datasets in **a.** different environments, **b.** in selected host datasets for which metadata detailing the host species were available and **c.** from different sampling sites of healthy human hosts for which detailed metadata on body sampling site were available. Numbers on the right indicate (*Listeria* positive samples/total samples) *per* category.

*Listeria* was detected in 14.05% 16S datasets (Fig. 1a). Note that no positive result could be obtained using our approach (see Materials and Methods) analysing full metagenomes, in line with the relative low abundance of *Listeria* species^32^ and consistent with a higher sensitivity of 16S sequencing compared to full metagenome sequencing for a given sequencing depth^33^. *Lm* was most frequently present in soil (673/1,700; mean relative abundance 1.2×10^-4^), sludge (70/309), sediment (32/170) and host-associated samples (854/7,695; mean relative abundance 9.0×10^-5^). Only few water (42/1,980) samples and no air sample (0/53) were positive for any *Listeria* species (Fig. 1a, and Fig. S1a for normalised data *per* category). *Lm* was the most prevalent *Listeria* species in both soil and host-associated environments (Fig. 1a). In samples where more than one *Listeria* species was present, *Lm* was significantly more abundant than other *Listeria* species, both in soil and hosts (Fig. 1a and Fig. S1b). We next investigated *Listeria* species host range (Fig. 1b). *Lm* was found to be the most abundant (mean relative abundance 5.5×10^-3^) and prevalent in cattle (80/1,270; 6.30%), which have indeed been reported as a potential reservoir for *Lm*^26^, especially hypervirulent clonal complexes^3,8,15^. We detected *Lm* in human samples at a similar prevalence than in cattle (173/3,338; 5.18%), but 40 times less abundantly (mean relative abundance 1.3×10^-4^). *Lm* was also frequently found in chicken (mean relative abundance 3.6×10^-4^, prevalence 28/552, 5.05%) and pig samples (mean relative abundance 4.7×10^-4^, prevalence 48/300, 16%) but not that of goats (0/212), where only *L. ivanovii* was detected, consistent with the known enrichment of *L. ivanovii* in small ruminants^34^. A high *Lm* prevalence in pigs and wild boars has been reported^35–37^, and pigs might constitute an underappreciated niche for *Lm*. We next investigated the human sampling sites in which *Lm* was present. As expected for a foodborne pathogen, *Lm* was detected in faecal samples (108/2,238), but also in oral (7/108) and sputum (3/50) samples (Fig. 1c), consistent with reports that *Lm* may colonise both the gut and the oral cavity^38,39^*. Lm* was rarely present in skin samples (2/56) and absent in vaginal samples (0/30), but for both categories only few datasets were available for analysis. The non-pathogenic species *L. innocua* and *L. seeligeri* were not detected in any human-associated samples, while *L. ivanovii*, the only other pathogenic *Listeria* species, was detected, albeit far less than *Lm*, second most frequently in human stools (Fig. 1c).

We aimed to replicate the result of frequent *Lm* carriage in human independently, and assessed *Lm* presence by *hly* PCR in the stools of a cohort of 900 healthy and a cohort of 125 diarrheic individuals (see Material and methods). It was detected in 10% (90/900) of healthy human stool samples and 20.8% (26/125) of diarrheic stools samples (Fig. S1c, Table S1). The enrichment of *Lm* in diarrhoea samples (χ^2^=11.702, *P*=0.0018, Benjamini-Hochberg correction) is consistent with the observation that *Lm* can induce diarrhoea^40,41^. The two-fold higher carriage level in healthy asymptomatic donors in this cohort from France, relative to the 16S datasets from MG-RAST may be due to the different sensitivities of the two methods (targeted *hly* amplification *versus* total 16S amplification), and sample selection bias reflecting a potential differential exposure to *Lm*-contaminated food^30^. Neither age nor gender was associated to asymptomatic carriage (Table S2).

The gut microbiota is a major line of defence against foodborne pathogens, and several commensals exert a protective effect against enteropathogens^42^, including *Lm*^43^. *Lm* also produces bacteriocins that can alter microbiota composition^44,45^. In order to assess if microbiota composition has an impact on *Lm* faecal carriage in human and *vice versa*, we investigated the relative abundance of microbiota taxonomic groups in MG-RAST human faecal samples. To take into account the compositional nature of data of different origins^46^, we calculated the ratios between microbiota phylogenetic groups and *Lm* abundance in the human microbiome datasets where *Lm* is present (Fig. 2, Supplementary Tables S3 and S4). *Lm* abundance correlated with the ratio of abundance of Firmicutes to Bacteroidetes phyla (Fig. 2a left), consistent with the observation that an increase of this ratio correlates with increased susceptibility to *Lm*^47^. This correlation is not due to *Lm* itself, as this species was excluded when the relative abundance of Firmicutes was calculated. The ratio of Actinobacteria to Bacteroidetes also correlated with *Lm* abundance (Fig. 2b left), and Actinobacteria were also significantly enriched compared to Firmicutes and Proteobacteria (Table S3). *Lm* abundance also correlated positively at the family and order levels with Lachnospiraceae (Fig. 2c left), Coriobacteriales (Fig. 2d left), Actinomycetaceae (Fig. 2e left), Erysipelotrichaceae (Fig. 2f left), and negatively with Porphyromonadaceae (Fig. 2g left). Erysipelotrichaceae have previously been reported to be elevated in asymptomatic *C. difficile* carriers, which suggests that loss of colonisation resistance is associated with this family^48^. In line with our results, a protective effect of Porphyromonadaceae has also been observed against *Salmonella enterica* serovar Typhimurium^49^, *Enterococcus faecium*^50^ and *C. difficile*^51^. The aforementioned significant associations with *Lm* abundance in faecal carriers were also found significant between carriers and non-carriers (Fig. 2, right panels), with the exception of Porphyromonadaceae for which only a trend was observed (Fig. 2f, right). For Lachnospiraceae, non-carriers showed a significantly higher prevalence than carriers (Fig. 2c, right), reflecting that comparisons between carriers and non-carriers are prone to study- and sample-dependent biases. Carriers also displayed less diverse microbiomes than non-carriers (Fig. S2a), a finding consistent with the observation that α-diversity is also involved in colonization resistance^42^ against enteropathogens such as *C. difficile*^52^*, Salmonella* or *Shigella*^53^. The overlap between the microbiota features associated with intestinal colonization by *Lm* and other well-known gut-colonising bacteria is consistent with our finding that *Lm* is frequently present in stools of asymptomatic individuals.

**Figure 2.**
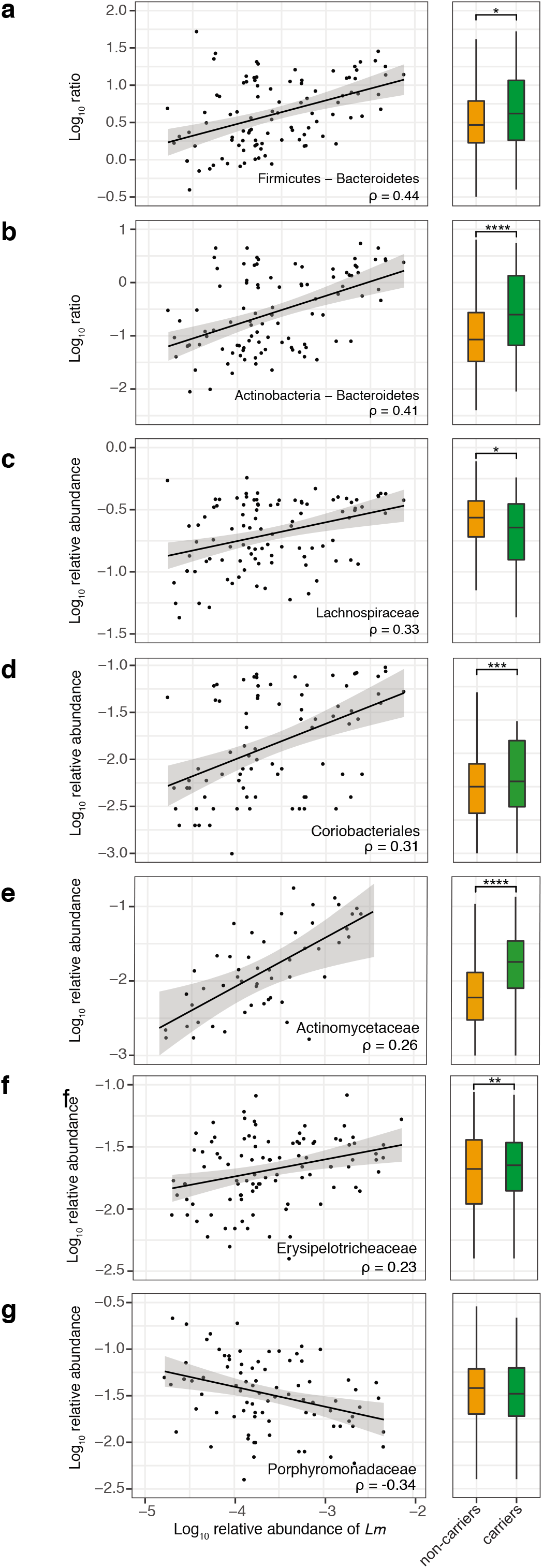
*Lm* carriage correlates with a specific microbiota signature in humans. All significant correlations with more than 75 associated samples and rho>0.2 between *Lm* and commensals relative abundance in 108 healthy carrier (left panels) and comparison between carriers and non-carriers for the same groups (right panels): **a.** The ratio of Firmicutes to Bacteroidetes phyla (rho=0.44, p=2.75×10^-5^, note that *Lm* species was excluded when the relative abundance of Firmicutes was calculated), **b.** The ratio of Actinobacteria to Bacteroidetes (rho=0.414, *P*=6.1×10^-5^), **c.** Lachnospiraceae (rho=0.326, *P*=1.25×10^-3^), **d.** Coriobacteriales (rho=0.314, *P*=4.01×10^-2^), **e.** Actinomycetaceae (rho=0.265, *P*=7.18×10^-11^), **f.** Erysipelotrichaceae (rho=0.226, *P*=4.51×10^-2^), **g.** Porphyromonadaceae (rho=−0.337, *P*=4.28×10^-3^). Statistical comparison performed with two-sided Wilcoxon rank-sum test with Benjamini-Hochberg correction for multiple test. * *P*<0.05, ** *P*<0.01, *** *P*<0.001 *\*\*\*\* P* <0.0001.

*Lm* shedding from infected tissues back in the intestinal lumen may favour long-term faecal carriage and account for the purifying selection of its virulence genes^6,9,10^, in line with the finding that the most virulent *Lm* clonal complexes are the most adapted to mammalian gut^8^, and the present observation that non-pathogenic species are not found in stool datasets retrieved from MG-RAST. To study *Lm* faecal carriage and its determinants experimentally, we inoculated mice intravenously with 5×10^3^ CFUs of *Lm* belonging to the hypervirulent clonal complex-1^8,54^. We observed a cage-dependent asymptomatic faecal carriage in 3/7 cages (11/26 mice). *Lm* could be detected over 30-days post-inoculation. We classified faecal carriage as either heavy (>10^6^ CFU/g, 6 mice in 2 cages) or light (<10^6^ CFU/g, 4 mice in 1 cage, together with one non-carrier mouse) (Fig. 3a). In 4 cages (15 mice), no *Lm* was detected in the faeces 30-days post-inoculation (Fig. 3a). All mice recovered from infection as assessed by weight gain, independently of their carrier status (Fig. S2b). We also separated mice and observed persistent faecal carriage, ruling out that it was resulting from coprophagy.

**Figure 3.**
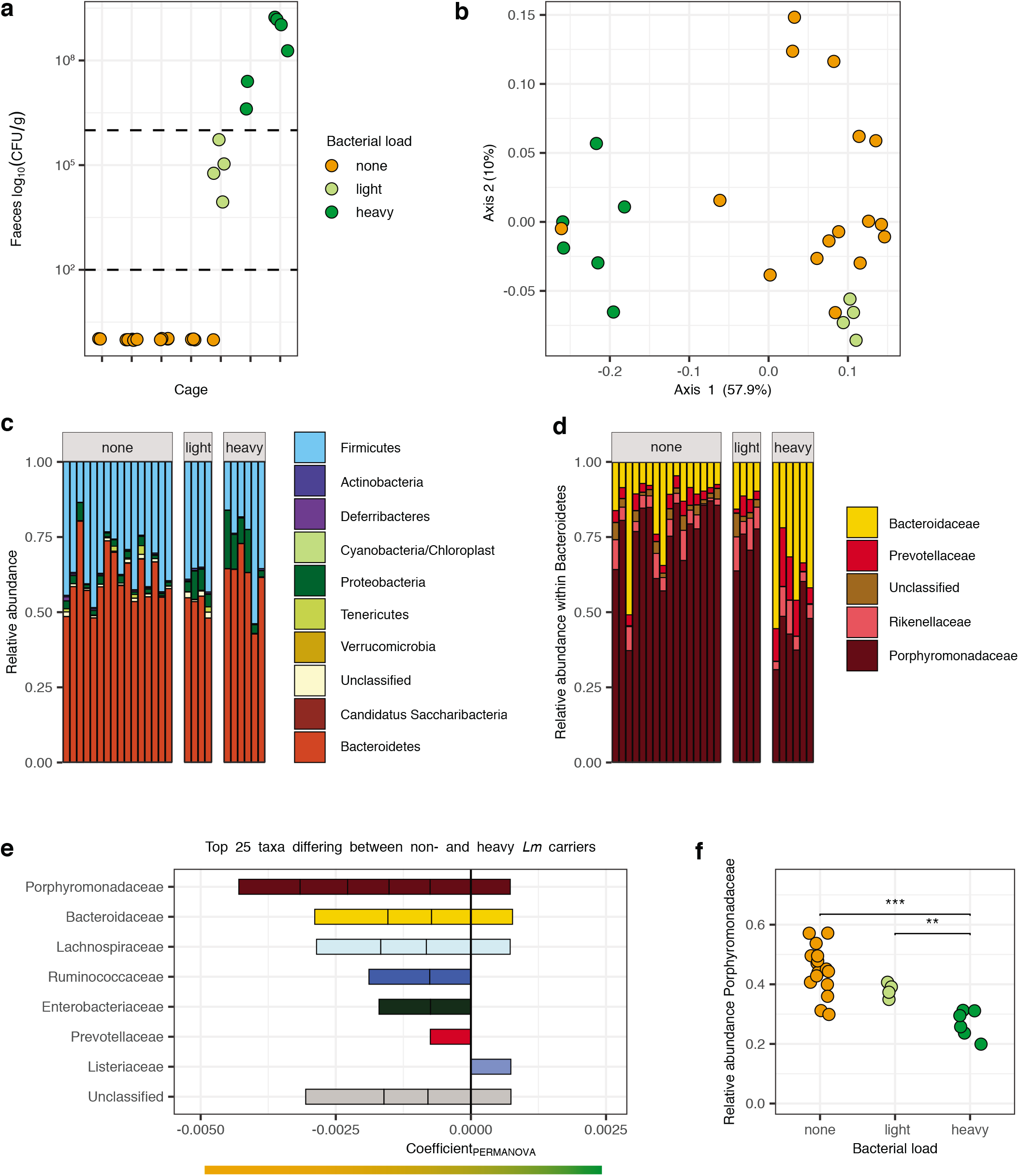
*Lm* long-term carriage correlates with a specific microbiota signature in mice a. CFU/g of stool of male mice 30 days after an *iv* challenge with *Lm* at 5×10^3^ CFU from different cages (2-6 mice per cage). Color indicates carriage group (<100 CFU/g: none, 100-10^6^ CFU/g: light, >10^6^ CFU/g: heavy). Horizontal lines indicate the threshold between the groups. **b.** β-diversity of mice microbiomes using MDS and Bray-Curtis distance. The color indicates the carriage group (<100 CFU/g: none, 100-10^6^ CFU/g: light, >10^6^ CFU/g: heavy). All groups differed in composition (PERMANOVA overall *P*=0.001, heavy/none *P*=0.006, heavy/light *P*=0.0075, light/none *P*=0.031, with Benjamini-Hochberg correction). Light carriers were more homogeneous than other groups (permutation test for homogeneity of multivariate dispersion, heavy/none, *P*=0.246, heavy/light *P*=0.0160, light/none *P*=0.0193, with Benjamini-Hochberg correction) **c**. Microbiota composition of mice from Fig. 2c at phyla level and **d.** family level within the Bacteroidetes phylum. **e.** PERMANOVA coefficients of 25 most different taxa between heavy and none-carriers microbiota from (Fig. 3b) according to their family. Horizontal bar indicates microbiota association (orange: none, green: heavy) **f.** Relative abundance of Porphyromonadaceae in 16S data from mice from different carriage groups. Statistical comparison performed with two-sided Wilcoxon rank-sum test. * *P*<0.05, ** *P*<0.01, *** *P*<0.001 *\*\*\*\* P*<0.0001.

Co-housed animals tend to have similar microbiota^55^, therefore the cage dependency of the observed differences in *Lm* carriage suggested that it was mediated by differences in gut microbiota composition. Indeed, heavy, light and non-carrier microbiota differed in microbial richness (α-diversity) and composition (β-diversity): heavy carriers’ microbiota was less diverse than that of light and non-carriers (Fig. S2c), consistent with results in human (Fig. S2a). β-diversity analysis showed that faecal carriage groups differed also in composition (PERMANOVA *P*<0.001): heavy carriers clustered separately from light and non-carriers (Fig. 3b). The difference between the light carrier group and the others reflected the higher homogeneity of the former (Fig. 3b), and the difference between non- and heavy carriers was mainly driven by a different composition in Bacteroidetes: 7 and 4 out of the 25 most contributing taxa belonged to Porphyromonadaceae and Bacteroidaceae, respectively (Fig. 3c-e). Indeed, Porphyromonadaceae were less abundant in the permissive microbiota of heavy carriers than that of light and non-carriers, and were present at an intermediary level in light carriers (Fig. 3f), similar to our observation in human (Fig. 2g).

We finally investigated whether these differences in microbiota α- and β-diversities result from or precede *Lm* carriage. We compared microbiota 16S composition before *Lm* inoculation and 30-days post-inoculation. *Lm* inoculation did not affect microbiota α-diversity (Fig. S3a). The β-diversity difference observed between heavy and non-carriers (Axis 1 in Fig. S3b) pre-existed *Lm* inoculation, and was mainly driven by a differential abundance of Porphyromonadacae (Fig. S3c and Fig. 3f). This pre-existing microbiota composition difference suggests that it plays a causative role in *Lm* carriage. *Lm* inoculation also had a significant impact on microbiota composition (Axis 2 in Fig. S3b). We investigated the nature of this microbiota change (Fig. S3d), and apart from the presence of *Lm* itself, we observed a decrease in Prevotellaceae upon *Lm* inoculation (Fig. S3e), likely reflecting the impact of *Lm* bacteriocin Lmo2776 on this bacterial family^44^.

Here we have shown that asymptomatic *Listeria* faecal carriage correlates with virulence: it is common in pathogenic *Listeria* species and absent in non-pathogenic species. Asymptomatic faecal carriage could thus be the force exerting purifying selection on virulence genes, rather than disease, which is rare and for which there is no inter-human transmission^56^. This also implies that humans are not a focal host for *Lm*. Consistent with this, *Lm* is more prevalent and abundant in cattle than in human stools, which is also in line with our recent report that hypervirulent *Lm* clonal complexes are associated to cattle and dairy products^8^. Moura *et al.* now report that the phylogeography of the hypervirulent *Lm* clonal complex-1 is linked to cattle global trade and farming^57^. Taken together, these observations strongly suggest that cattle constitute a major reservoir where *Lm* virulence is selected for.

We also found that *Lm* is the predominant *Listeria* species in the environment, where it is a saprophyte. *Lm* persistence in food processing plants, away from its natural hosts, is associated with loss of virulence^7,8,54^. That *Lm* is found more abundantly in soil, sludge and sediments than non-pathogenic species (Fig. S1a) suggests that *Lm* regularly transits between its hosts *via* these environments, while maintaining its host-association capacity, which is mediated by its “virulence” genes. *Listeria* host-association capacity therefore appears as a trait that ensures the ecological success of *Lm* and *L. ivanovii* relative to other species. The relative lower prevalence in the environment of non-virulent *Listeria* species *L. innocua*, *L. seeligeri* and *L. welshimeri* which derive from the common virulent ancestor of *Lm* and *L. ivanovii*^58^ suggest that they either (*i*) successfully colonize an environment not sampled in this study, and/or (*ii*) lost their focal host, and/or (*iii*) lost their host association capacity, similar to *Lm* clones associated with food processing plants which are in the process of losing virulence^7,8,54^. It will be interesting to investigate how host association is also involved in the overall ecological success of other microbial species which, as *Lm,* are widespread in the environment. Future research will also have to address the relative contribution of host and *Lm* genetics, food habits, and intestinal microbiota to asymptomatic faecal carriage of *Lm*.

## Acknowledgments

We thank Georges Michel Haustant and Cédric Fund, Biomics Platform, C2RT, Institut Pasteur, Paris, France, supported by France Génomique (ANR-10-INBS-09-09) and IBISA for 16S sequencing, Auguste Fourneau and Amandine Brunet for PCR assays, and Sandrine Isaac, Henrik Salje and Olivier Disson for critical reading. LH is supported by the Pasteur-Paris University (PPU) International PhD Program, funded by the European Union’s Horizon 2020 research and innovation programme under the Marie Sklodowska-Curie grant agreement No 665807, and the “Ecole Doctorale FIRE-Programme Bettencourt” of the CRI Paris. This work was funded by Institut Pasteur, Inserm, Laboratoire d’Excellence Integrative Biology of Emerging Infectious Diseases and the European Research Council.

## Author contributions

ML initiated and coordinated the project. MGG, LH, AM and ML designed the study. MGG collected and analyzed the public metagenomic data. CB and MP assessed the prevalence of *Lm* in stool donors’ cohorts. LH conducted the *in vivo* experiments in mice and the corresponding 16S rDNA analysis. LH and ML wrote the manuscript, MGG and AM commented and edited on it.

## Materials and methods

### Screening of Listeria sp. in 16S rDNA datasets

A summary of the study workflow is represented in Fig. S4. We collected 13,749 16S rDNA amplification datasets from MG-RAST from studies with >5 and <50 samples, for studies containing host samples <250 (last accessed: November 2017) as described in ^59^. When more samples were available, we randomly selected 50 or 250 samples, respectively. We removed those containing non-ribosomal data or less than 2000 sequences using SSU-align v.1.01^60^. This left us with a total of 11,907 rDNA datasets (Supplementary Table S5). Sequences shorter than 60 bp were removed. 16S rDNA sequence datasets were re-aligned using mafft v. 7.407^61^, and trimmed using trimal v.1.4^62^ using the ‘automated1’ algorithm. The resulting trimmed sequences were then clustered within each sample at 99% identity and 90% coverage using the uclust algorithm from usearch v. 10.0.240^63^. A representative sequence of each cluster was defined according to the distance to the cluster centroid. Henceforth, we will call these our environmental dataset.

To identify *Listeria* ssp. in the environmental dataset we used a maximum likelihood approach. First, *Lm* 16S rDNA reference sequences were aligned using mafft with the ‘linsi’ algorithm. The resulting multiple sequence alignment was trimmed using trimal v.1.4. A phylogenetic reconstruction was then performed using IQ-tree v.1.6.5^64^ using the GTR model (according to the model test) and 1000 rapid bootstrap iterations. The resulting tree was manually pruned to leave only one representative member of each clade. Environmental sequences were then classified as potential *Listeria* candidates by mapping them against the multiple sequence alignment using the ‘-addfragments’ algorithm of mafft. Sequences with at least 90% identity and 90% coverage to one reference member were kept for further analyses, or otherwise were discarded. The remaining sequences were then assigned to one of the branches of the phylogenetic tree using the evolutionary placement algorithm implemented in RAxML v. 8.2^65^.

Environmental sequences assigned to any terminal branches with a maximum likelihood of 0.6 or higher, were classified as the specific *Listeria* species. Otherwise, they were classified as “*Listeria* undefined”. Note that this was the case for sequences with a non-discriminative amplicon region at the species level, e.g. V3-V4. In this work, we focused on all *Listeria sensu stricto* species, which are frequently found in the environment (*Lm*, *L. ivanovii*, *L. innocua*, *L. seeligeri* and *L. welshimeri*). We did not include the closest non-pathogenic relative of *Lm*, *L. marthii* since it is only rarely sampled in any environment^66^.

The remaining representative sequences were used to construct a global catalogue of operational taxonomic units (OTUs). To do so, the representative sequences of all datasets were grouped and clustered together at 97% identity using usearch, and the frequency of each OTU was calculated on each dataset. Finally, OTU representatives were taxonomically classified at genus level using the RDP classifier^67^. At the same time, we defined the α-diversity of each dataset as the Expected Number of Species (ENS). To do so, we did calculate the Shannon diversity index (H’):

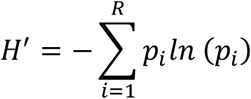

where p_i_ is the relative frequency of a specific species in the dataset (the number of sequences associated with the species divided by the total number of sequences assigned to species), and R is the number of datasets. We calculated the ENS as the exonential of the Shannon diversity:

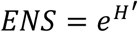

### Screening of Listeria sp. in 16S full metagenomes

We followed the approach described in Garcia-Garcera et al 2017. In brief, we retrieved around 3,000 metagenomes from MG-RAST. To characterize the presence of *Lm*, reads were mapped against a discriminative concatenate of genes, here the 7 housekeeping genes used for multilocus sequence typing in *Lm*^68^. Since no reads could be mapped to these gene concatenate, no further downstream analysis has been performed. The absence of Lm in full metagenomes can either be due to the low relative abundance of Lm per se which does not permit it to be identified, or previous filtering of reads from species with low abundance.

### Detection of Lm in human faecal samples

Determination of faecal carriage of *Lm* was performed by PCR amplifying *hly*^24^. We evaluated the performance of this method using artificial samples that mimic natural stool. Briefly, 10-fold dilutions of the ATCC *Lm* strain (ATCC BAA751) in saline buffer (10^8^ to 10^1^ CFU/mL) were diluted in a 1:1 ratio in a PCR-negative stool sample conserved on eNat (Copan, Italy) before extraction. Extraction was performed with EasyMag (bioMérieux, Marcy-l’Etoile, France) according to manufacturer’s recommendations. PCR assays were performed in triplicate on a CFX96 system (BioRad, CA, USA) as described ^24^. *Lm* was considered present when at least 2 of 3 PCR assays were positive. *Lm* detection threshold was 10^6^ CFU/ml of stool. Tested stool samples originated from two cohorts collected and stored on eNat (Copan): (*i*) the Hepystool cohort that included samples (n=900 samples, 2015-2016) from non-diarrheic patients (inclusion criteria described in ^69^) and (*ii*) stool samples from diarrheic patients, received at the Infectious Agents Department of the University Hospital of Poitiers, France. DNA was extracted on EasyMag (bioMérieux, Marcy-l’Etoile, France) according to manufacturer’s recommendations then amplified in triplicate. All samples which were at least once positive on the first triplicate were subjected to a second triplicate and were considered as positive when again detected at least once.

Note that no culture-based identification was applicable given that the eNat protocol conserves nucleic acid and is bactericidal within 30 minutes (see manufacturer’s instructions for details).

### Mouse colonization experiments

7 to 11-week-old male mice (C57BL/6 mEcad E16P KI^70^) were infected intravenously in the tail vein as previously described^70^. Fraternities were kept together in a cage during the whole experiment, except when separated to exclude that carriage was due to coprophagy. To quantify carriage at 30-days post-inoculation, faeces were collected from each individual mouse and weighted before being homogenized in 2ml of PBS. CFU count was performed by serial dilution of homogenized faeces on ALOA plates as described in ^8^. Separate faecal pellets were collected pre-inoculation and/or 30 days-post-inoculation and stored at −20°C for DNA extraction for 16S sequencing.

### 16S rDNA analysis in mice

DNA from faeces has been isolated with DNeasy PowerSoil Kit (Qiagen) accordingly to the manufacturer’s instructions. The V4 region has been amplified and sequenced with the primers CCTACGGGNGGCWGCAG and GACTACNVGGGTWTCTAATCC using the Illumina MiSeq workflow at the biomics platform at the Institut Pasteur, Paris. Analysis have been performed with micca^71^, using the RDP classifier^72^ and unoise3 for clustering^73^. Forward and reverse reads were merged with a minimum overlap of 100bp and 30 maximum allowed mismatches. Forward and reverse primers were removed and reads were trimmed to 400 nucleotides using the mica workflow. Reads with an expected error rate above 0.75% were excluded. Reads were grouped in sequence variants by unoise3^73^ and chimeric sequences were removed. Sequence variants were classified with RDP^72^, which uses VSEARCH to match sequences with the reference database^74^. Statistical were performed with R and the phyloseq, vegan and microbiome libraries^75,76^. α-diversity has been calculated by number of observed species, abundance-based coverage estimator (ACE)^77^ and Shannon index^78^. β-diversity between samples has been calculated with MDS of Bray-Curtis dissimilarities^79^. PERMANOVA and homogeneity between microbiome groups were calculated with adonis and betadispers from the vegan library^80^.

### Ethical statement

Animal experiments were performed according to the Institut Pasteur guidelines for laboratory animals’ husbandry and in compliance with European regulation 2010/63 EU. All procedures were approved by the Animal Ethics Committee of Institut Pasteur, authorized by the French Ministry of Research and registered under #11995-201703115103592 and #14644- 2018041116183944. The stool donor cohorts received ethical approval from the regional Committee for the Protection of People (CPP Ouest III) and from the National commission for Protection of Personal data on November 23^th^ 2015. All patients were informed before inclusion and their consent was obtained before analysis.

## Data availability

Primary sequencing data are available on the Sequence Read Archive under the entry PRJNA642013

---

**Supplementary Figure 1.**
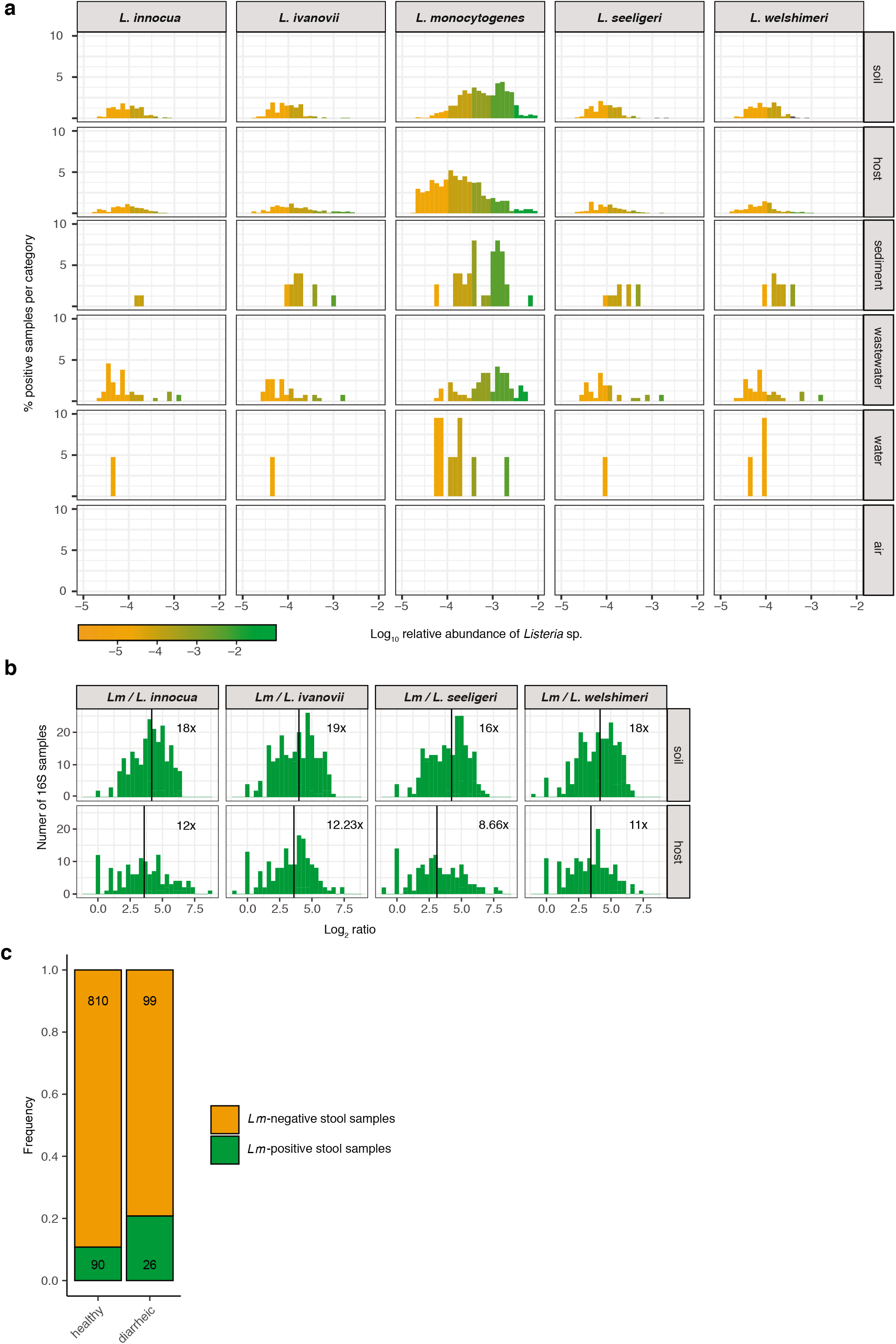
*Lm* is more abundant than other *Listeria* species in all environments and *Lm* carriage is common in healthy individuals. (Related to Figure 1) **a.** Same as Fig. 1a normalised by category. **b.** Log2 of ratio of *Lm* to each other evaluated *Listeria* species in samples where the species co-occurred. Vertical line and number indicate the mean of the distribution. **c.** Prevalence of *Lm* in human faecal samples from healthy (n=900) and diarrheic donors (n=125) from France.

**Supplementary Figure 2.**
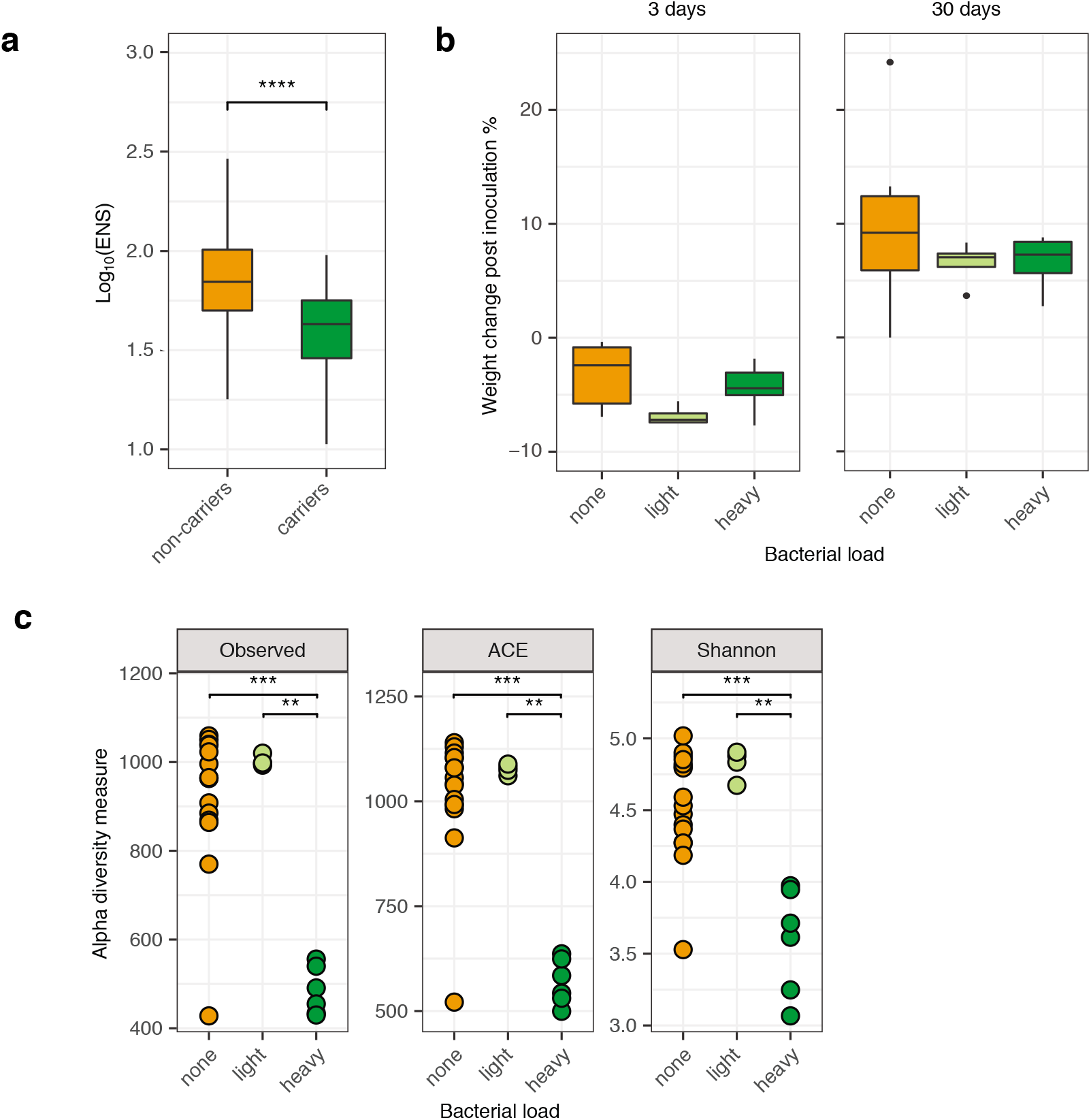
*Lm* carriage correlates with low α-diversity. (Related to Figures 2 and 3) **a.** α-diversity, measured by ENS between carriers and non-carriers, similar to Fig. 2B. **b.** Body weight change of mice after inoculation at 3 days post-inoculation and 30 days post-inoculation according to their carriage group. **c.** Carriage groups differ in α-diversity, measured by observed species (left), abundance-based coverage estimate (middle) and Shannon index (right). Statistical comparison performed with two-sided Wilcoxon rank-sum test. * *P*<0.05, ** *P*<0.01, *** *P*<0.001 *\*\*\*\* P*<0.0001.

**Supplementary Figure 3.**
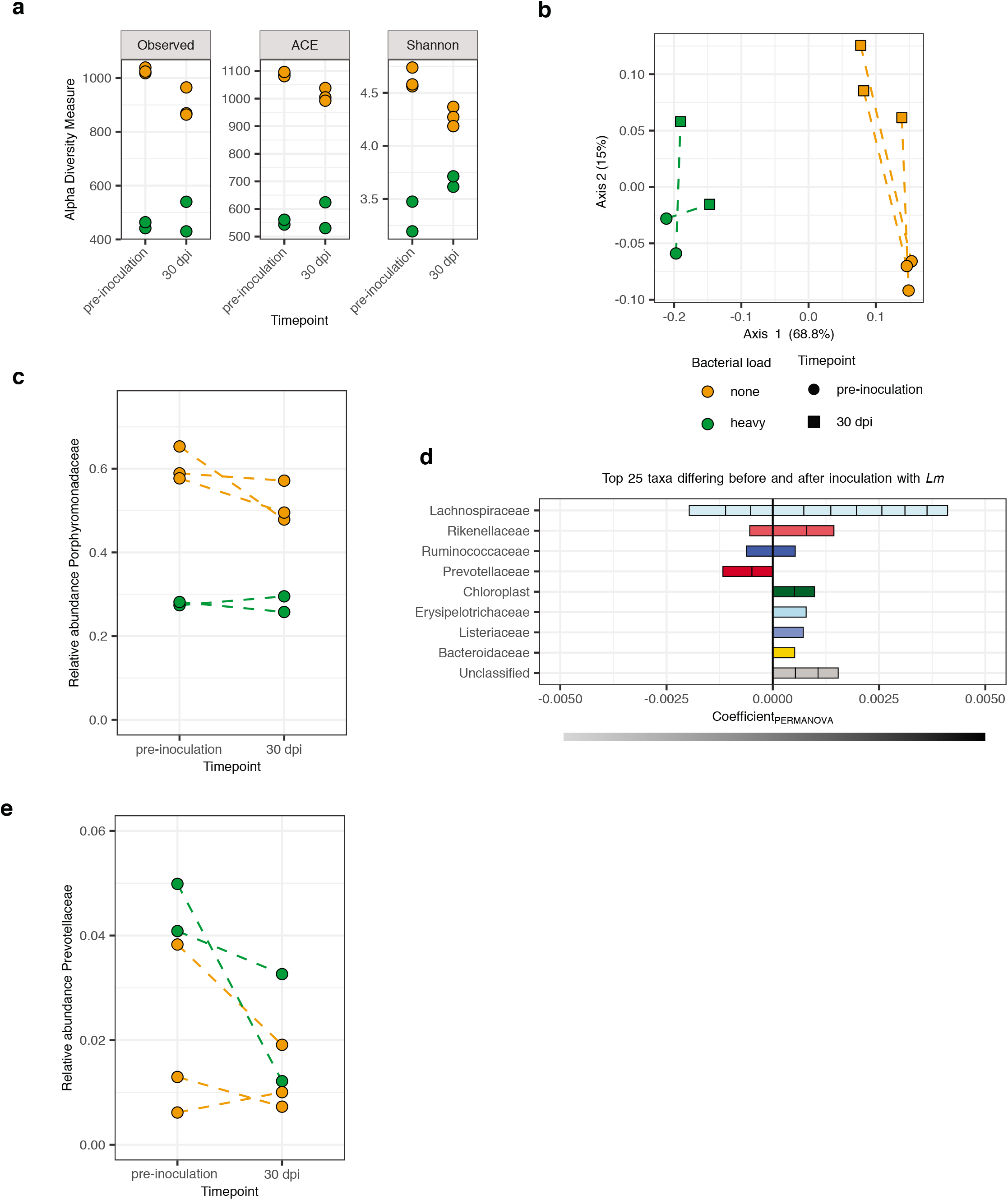
Effect of *Lm* inoculation on microbiome composition. (Related to Figure 3) Comparison of microbiota pre-inoculation and 30-days post-inoculation for 2 cages with 3 non-carrier and 2 heavy carrier mice. **a.** α-diversity, measured by observed species (left), abundance-based coverage estimates (middle) and Shannon index (right) before inoculation and 30-days post-inoculation. Statistical comparison performed with two-sided Wilcoxon rank-sum test. **b.** β-diversity of mice microbiomes using MDS and Bray-Curtis distance. The colour indicates the carriage group (<100 CFU/g: none, >10^6^ CFU/g: heavy) and the shape the timepoint (round: Pre-inoculation, square: 30-days post-inoculation). **c.** Relative abundance of Porphyromonadacea pre-inoculation and 30-days post-inoculation. **d.** PERMANOVA coefficients of 25 most different taxa between microbiomes pre-inoculation and 30-days post-inoculation (Fig. S3b). Horizontal bar indicates microbiota association (grey: pre-inoculation, black: post-inoculation) **e.** Relative abundance of Prevotellaceae pre-inoculation and 30-days post-inoculation.

**Supplementary Figure 4.**
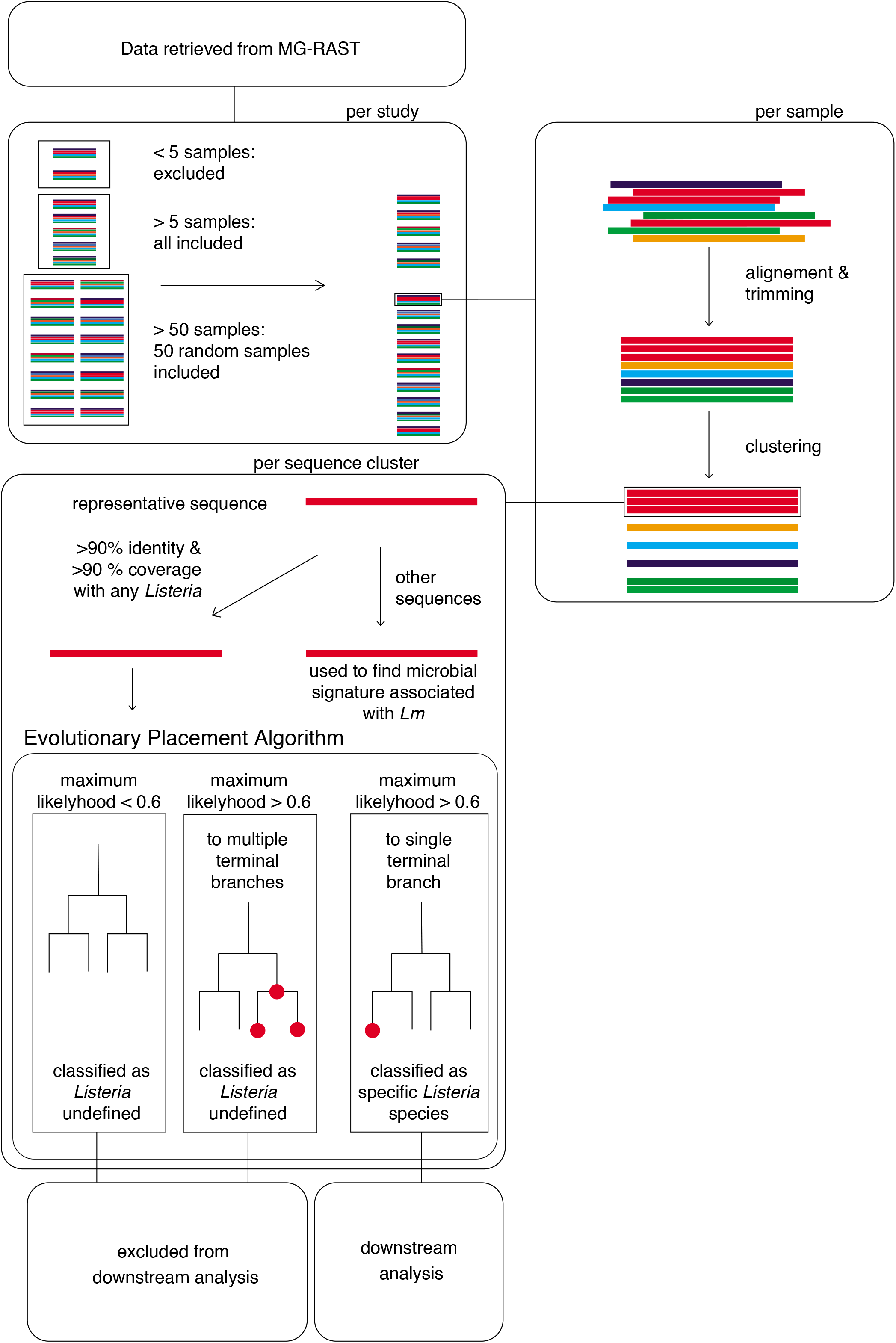
Workflow of screening of *Listeria sp.* in 16S rDNA datasets. (Related to Material and Methods) Data was retrieved from MG-RAST. Samples from studies with <5 samples were excluded, samples from studies with >5 and <50 samples and random selections of samples from studies with >50 samples were included. For each sample, sequences were aligned and trimmed, and then clustered. A representative sequence from each sequence cluster was then mapped to *Listeria sp.* sequences. Sequences with >90% identity and coverage with at least one *Listeria* were assigned to a tree of *Listeria* reference sequences by a evolutionary placement algorithm. Sequences with a maximum likelihood >0.6 for a single terminal branch were assigned to the corresponding species, others were excluded from downstream analysis.

## Supplementary Tables

**Supplementary Table S1.** Sensitivity of *hly* PCR in different matrices

**Supplementary Table S2.** Metadata of stool collection cohort

**Supplementary Table S3.** Correlations of *Lm* abundance with microbial phyla

**Supplementary Table S4.** Correlations of *Lm* abundance with microbial families and orders

**Supplementary Table S5.** List of 16S datasets from MG-RAST used in this study

