## Supplementary material for "*Listeria monocytogenes* faecal carriage is common and driven by microbiota": Tables S1 and S2

Supplementary Table 1: Sensitivity of *hly* PCR in different matrices

| | -Stool sample | | +Stool sample | |
|---|---|---|---|---|
| CFU | SB | EB | FSB | FEB |
| $1 \times 10^8$ | 3/3 | 3/3 | 3/3 | 3/3 |
| $1 \times 10^7$ | 3/3 | 3/3 | 2/3 | 2/3 |
| $1 \times 10^6$ | 3/3 | 3/3 | 2/3 | 2/3 |
| $1 \times 10^5$ | 3/3 | 3/3 | 3/3 | 1/3 |
| $1 \times 10^4$ | 3/3 | 2/3 | 1/3 | 0/3 |
| $1 \times 10^3$ | 3/3 | 0/3 | 1/3 | 0/3 |
| $1 \times 10^2$ | 3/3 | 0/3 | 0/3 | 1/3 |
| $1 \times 10^1$ | 0/3 | 0/3 | 0/3 | 0/3 |

SB: Saline buffer, EB: eNat buffer, FSB: Negative stool sample diluted in saline buffer, FEB: Negative stool sample diluted in eNat buffer. The indicated amounts of *Lm* were resuspended in each matrix and a PCR of the *hly* gene was performed three times per sample. Red indicates samples that would be considered as negative with less than two positive PCR results, green indicates samples that would be considered as positive with at least two positive PCR results.

Supplementary Table 2: Metadata of stool collection cohort

| | *Lm*-positive (n=90) | *Lm*-negative (n=810) | *P*-value |
|---|---|---|---|
| Median age (year; [IQR]) | 47.7 [38.2-60.7] | 52.5 [39.9-63.6] | 0.1812 |
| Median delay before reception of the stool (day; [IQR]) | 1.00 (1-2) | 1.00 (1-2) | 0.5255 |
| Male, n (%) | 33 (36.7) | 340 (42.0) | 0.3677 |
| Female, n (%) | 57 (63.3) | 470 (58.0) | |
| Birthplace | | | |
| Europe | 85 (10.4) | 730 (89.6) | 0.2524 |
| Africa | 3 (1.4) | 66 (98.6) | 0.1407 |
| Asia | 0 (0) | 8(100) | 1.00 |
| America | 2 (25) | 6 (75) | 0.2079 |
